# Adaptation, Clonal Interference, and Frequency-Dependent Interactions in a Long-Term Evolution Experiment with *Escherichia coli*

**DOI:** 10.1101/017020

**Authors:** Rohan Maddamsetti, Richard E. Lenski, Jeffrey E. Barrick

## Abstract

Twelve replicate populations of *Escherichia coli* have been evolving in the laboratory for more than 25 years and 60,000 generations. We analyzed bacteria from whole-population samples frozen every 500 generations through 20,000 generations for one well-studied population, called Ara-1. By tracking 42 known mutations in these samples, we reconstructed the history of this population’s genotypic evolution over this period. The evolutionary dynamics of Ara-1 show strong evidence of selective sweeps as well as clonal interference between competing lineages bearing different beneficial mutations. In some cases, sets of several mutations approached fixation simultaneously, often conveying no information about their order of origination; we present several possible explanations for the existence of these mutational cohorts. Against a backdrop of rapid selective sweeps both earlier and later, we found that two clades coexisted for over 6000 generations before one drove the other extinct. In that time, at least nine mutations arose in the clade that prevailed. We found evidence that the clades evolved a frequency-dependent interaction, which prevented the competitive exclusion of either clade, but which eventually collapsed as beneficial mutations accumulated in the clade that prevailed. Clonal interference and frequency dependence can occur even in the simplest microbial populations. Furthermore, frequency dependence may generate dynamics that extend the period of coexistence that would otherwise be sustained by clonal interference alone.

## INTRODUCTION

The long-term evolution experiment (LTEE) spans more than 25 years and 60,000 generations of bacterial evolution. In this experiment, 12 replicate populations of *Escherichia coli* have been propagated in a simple environment, and samples of each population frozen at 500-generation intervals. This experiment originally focused on whether and to what extent the populations would diverge in their mean fitness and other phenotypic properties as they adapted to identical environments (Lenski *et al.* 1991; Lenski and Travisano 1994). Over time, this experiment has become a model for exploring many other aspects of evolution, including the emergence of new functions (Blount *et al.* 2008), the evolution of mutation rates (Sniegowski *et al.* 1997), the maintenance of genetic diversity (Elena and Lenski 1997; Rozen and Lenski 2000; Le Gac *et al.* 2012), and the structure of the fitness landscape (Woods *et al.* 2011; Khan *et al.* 2012; Wiser *et al.* 2013). The ability to examine these and other issues has grown tremendously as data that were difficult or impossible to obtain when the LTEE began have yielded to new technologies, particularly genome sequencing (Barrick and Lenski 2009; Barrick *et al.* 2009; Blount *et al.* 2012; Barrick and Lenski 2013; Wielgoss *et al.* 2013).

The LTEE has also inspired theoretical work, especially on the dynamics of adaptation in large asexual populations (Gerrish and Lenski 1998; Hegreness *et al.* 2006; Desai and Fisher 2007; Schiffels *et al.* 2011; Park and Krug 2013; Wiser *et al.* 2013). The LTEE populations are subject to clonal interference, a phenomenon that limits the rate of adaptation by natural selection in large asexual populations. In the absence of recombination, two or more beneficial mutations that appear in different lineages in the same population cannot recombine into a single background; instead, the lineages that possess alternative beneficial mutations compete with one another. As a consequence, each beneficial mutation will interfere with the progress of other contending beneficial mutations toward fixation, though the mean fitness of the population will nonetheless rise as the beneficial alleles collectively displace their progenitors. Although some early theory on clonal interference was developed with the LTEE in mind (Gerrish and Lenski 1998), other evolution experiments using bacteria and yeast have provided compelling demonstrations of this phenomenon by combining dense temporal sampling with intensive discrimination using genetic markers (De Visser and Rozen 2006; Hegreness *et al.* 2006; Woods *et al.* 2011; Barroso-Batista *et al.* 2014), in-depth analysis of genes under positive selection (Lee and Marx 2013), or whole-genome sequencing (Lang *et al.* 2011; Lang *et al.* 2013).

Without clonal interference, the classic model of periodic selection in asexual populations involves selective sweeps of beneficial mutations that arise singly and fix sequentially. Neutral and nearly neutral mutations that would otherwise accumulate in an evolving population are swept out (or occasionally to fixation) as each successive beneficial mutation goes to fixation (Atwood *et al.* 1951). As a consequence, within-population genetic diversity rises and falls in conjunction with the successive sweeps, and no specific polymorphism is maintained indefinitely. Clonal interference can increase genetic diversity in three ways. First, the multiple beneficial mutations, all rising to moderate frequencies, increase diversity relative to a single allele rising to high frequency. Second, the beneficial alleles remain at intermediate frequencies longer than would a single beneficial mutation of comparable effect-size during a classic sweep. Third, while the beneficial mutations remain at intermediate frequencies, their associated lineages can accumulate neutral and nearly neutral polymorphisms that also persist longer than they would in the face of selective sweeps that progress to fixation. Nonetheless, the diversity-promoting effects of clonal interference are only transient because eventually one lineage or another will prevail, either because the beneficial mutation it carries is superior to the others (even though all lineages are more fit than their predecessors) or because one lineage acquires additional beneficial mutations that eventually break the logjam.

Although clonal interference can prolong polymorphic states only transiently, ecological interactions between genetically diverged individuals in a population that result in negative frequency-dependent effects on fitness can sustain polymorphisms indefinitely, at least in principle. The LTEE environment was designed to minimize the potential for frequency-dependent interactions to arise, in order to simplify measuring fitness and assessing the repeatability of evolution. In particular, there are no predators, parasites, or other competing species in the LTEE environment; there is no sexual recombination or horizontal gene transfer; the physical environment is well mixed and lacks spatial structure; and the density-limiting resource, glucose, is provided at a low concentration, which limits the concentration of metabolic byproducts that could support cross-feeding specialists. Nonetheless, frequency-dependent interactions have emerged in some of the LTEE populations (Elena and Lenski 1997; Rozen and Lenski 2000; Rozen *et al.* 2009; Le Gac *et al.* 2012; Plucain *et al.* 2014; Ribeck and Lenski 2015).

In previous work, we sequenced individual clones and population samples taken at generations 2000, 5000, 10,000, 15,000, 20,000, 30,000, and 40,000 from one LTEE population, designated Ara-1, that has served as the focal population for many in-depth analyses (Barrick and Lenski 2009; Barrick *et al.* 2009). Using these genomic data, we designed genotyping assays for many derived alleles present in one or more clones or population samples. In this study, we use population samples taken at 500-generation intervals to examine at high resolution the dynamics of 42 mutant alleles over the first 20,000generations of this population. These data show compelling evidence for both adaptive fixations and clonal interference. The data also suggest, and competition assays confirm, that a previously unknown negative frequency-dependent interaction evolved that delayed the fixation of some mutations for several thousand generations—in effect extending the duration of a particular bout of clonal interference. Nevertheless, one clade’s slight advantage when rare was eventually overcome by the consolidation of further beneficial mutations in the other clade, and this bubble of transient diversity collapsed.

## MATERIALS AND METHODS

### Ara-1 Population Samples

The LTEE is described in detail elsewhere (Lenski *et al.* 1991; Lenski and Travisano 1994; Lenski 2004). In brief, 12 replicate populations have been propagated in a glucose-limited medium called DM25. Daily 100-fold dilutions and re-growth allow ~6.6 generations per day. Every 500 generations (75 days), after the transfers into fresh medium, glycerol was added as a cryoprotectant to the remaining cultures, which were then stored for later research at -80°C. In this study, we analyzed the 40 samples of population Ara-1 collected from 500 to 20,000 generations.

### Sample Preparation for Genotypic Analyses

We briefly thawed the top portion of each frozen sample and removed 0.1 ml. We washed the cells in a saline solution and centrifuged them to remove residual glycerol; we then inoculated the cells into flasks containing 10 ml of DM100 medium (the same medium used in the LTEE, except with a higher glucose concentration to yield more cells,) which were incubated for 24 h at 37°C. We also mixed fully-grown cultures of the Ara^+^ ancestral strain, REL607, and a 20,000-generation clone from population Ara-1, REL8593A, at several cell ratios for use as standards for calibrating the genotypic analyses. We then isolated genomic DNA from all of these cultures using an Invitrogen PureLink *Pro* 96 Genomic DNA Purification kit.

In a separate experiment, we revived the frozen samples from generations 7500 and 10,000using the same protocol as above. The next day, we diluted and plated samples from the cultures, and the following day we picked 90 clones (single-colony isolates) from each sample. We inoculated each of these clones into a flask containing 10 ml of DM1000 medium and isolated genomic DNA from these cultures after growth, as described for the population samples.

We quantified genomic DNA concentrations for all samples using an Invitrogen Quant-iT PicoGreen dsDNA Assay kit, and we then submitted the samples for genotyping to the MSU Genomics Core facility. We prepared and submitted three different biological replicates for the population-level analyses, each of which was separately revived from the frozen sample and re-cultured before DNA isolation.

### Allele Frequency Measurements

We performed an Illumina GoldenGate Genotyping Assay with Veracode technology to measure allele frequencies in the Ara-1 population. We designed allele-specific oligonucleotides for single nucleotide polymorphisms (SNPs), insertions, deletions, and rearrangements that were previously discovered by sequencing either clones or whole-population samples (Barrick and Lenski 2009; Barrick et al. 2009). These oligonucleotides contain universal PCR primer sequences and barcode sequences targeting specific Veracode microbead types. Allele-specific, fluorescent-labeled PCR products hybridize to the microbeads, and the fluorescent signal provides an indicator of allele frequency. Of the assays we designed, 42 SNPs and indels yielded useful data about the history of the Ara-1 population. The mutations are listed in Table S1.

Illumina’s GenomeStudio software gave initial estimates of allele frequencies. We used the allele frequency estimates generated from known mixtures of REL8593A and REL607 to correct the frequencies of 25 of the 42 mutations that were present in REL8593A (all those that had fixed in the population by 20,000 generations). These DNA samples consisted of 21 known mixtures of the two strains designed to contain every 5% increment, based on culture volumes, in the percentage of REL8593A from 0% to 100%. We measured the true ratio of cell numbers in each mixed sample by plating a dilution on tetrazolium arabinose (TA) indicator agar and counting the red and white colonies made by REL8593A and REL607, respectively.

Average cell size increased over time in the LTEE (Lenski and Travisano 1994), and the evolved bacteria also contain more DNA per cell than the ancestral strain (Lenski *et al.* 1998). This change in DNA content meant that the ratio of REL8593A to REL607 cells in the control mixtures had to be corrected to reflect the actual number of copies of the evolved and ancestral alleles in these samples. To account for this difference, we stained stationary - phase cultures of the ancestral Ara~ clone, REL606, and the 20,000-generation clone, REL8593A, with a PicoGreen fluorescent DNA stain (Ferullo *et al.* 2009) and used flow cytometry to quantify the average genomic DNA content per cell in DM100 media (Figure S3). We used six replicate measurements to calculate a correction factor for the allele frequencies in the REL8593A/REL607 mixtures. On average, the evolved REL8593A cells had 1.67 ± 0.23 (± 95% confidence interval) times as much DNA as the ancestral REL606 cells under these conditions.

With this information, we further corrected the raw frequencies estimated for each allele (θ_0_) from the GenomeStudio genotyping software to the known frequencies of that allele (θ) in the control DNA samples. From the triplicate assays of the REL8593A/REL607 mixtures, we fit a calibration curve for each allele to an empirical function that corrected for a symmetric convex bias that captured the deviation from linearity (Figure S4). Specifically, we fit five coefficients (*C*_n_) in a linear model of the form θ = θ_0_ + *C*_1_(θ_0_ - θ^2^) + *C*_2_(θ_0_ - θ_0_^4^) + *C*_3_(θ_0_ - θ_0_^4^) + *C*_5_(θ_0_ - θ_0_^8^) + *C*_5_(θ_0_ - θ_0_^10^) to each calibration curve. Then, we used these curves to correct the GenomeStudio estimates of allele frequencies in all of the Ara-1 population samples that we analyzed. These calculations, as well as the code used to plot the temporal dynamics of the allele frequencies, are available as R analysis scripts at http://www.github.com/rohanmaddamsetti/Ara-l_Genotyping.

For the 17 transient alleles we analyzed that were not present in the REL8593A genome, we could not correct their estimated allele frequencies in this way. These allele estimates are thus expected to be slightly less accurate (see Figure S4 for an example of the typical magnitude of this correction). In cases where these unsuccessful mutations were competing with a contemporary allele that fixed by 20,000 generations, we constrained their frequencies to being no more than 100% minus the prevalence of the most abundant successful mutation. In any case, the estimated frequency trajectory for each individual evolved allele is shown in Figure S1, while the Muller plot in Figure 1 is based on a simple interpretation of how the estimated frequencies of each genotype sum to 100%.

**Figure 1.**
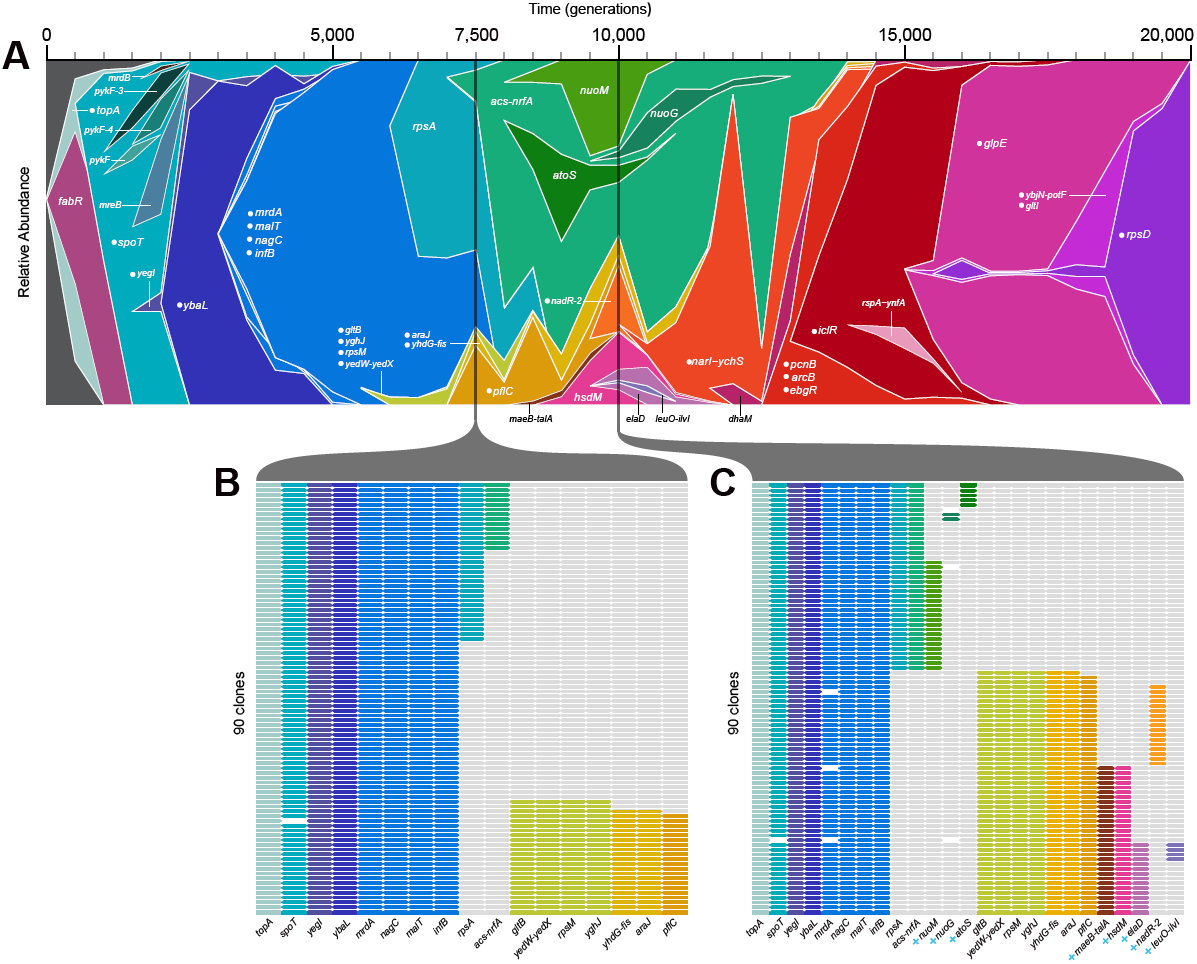
Dynamics of mutant alleles during a long-term evolution experiment with *E. coli.* (A) The Muller plot shows estimated frequencies of 42 known mutations in population Ara-1 over the course of 20,000 generations. Mutations are identified by the gene in which they occurred or, if intergenic, by the adjacent genes; trailing numbers indicate that the same gene had multiple alleles. Labels preceded by a dot indicate the mutation fixed in the population, as indicated by its presence on the line of descent leading to sequenced clones from 30,000 and 40,000 generations. Genotypes of 90 clones sampled at (B) 7500 and (C) 10,000 generations, showing previously fixed and variable alleles only. Each row in each panel represents a clone. Mutations are colored as in (A); light grey fill shows the mutation is not present (i.e., the clone has the ancestral allele), and white fill indicates missing data. Mutations labeled with the blue + symbol were detected in the clones sampled at 10,000 generations but not at 7500 generations.

### Pyrosequencing

We confirmed the existence of large temporal fluctuations in the two coexisting clades in the Ara-1 population by using pyrosequencing to estimate the *rpsA, yghj,* and *gltB* allele frequencies in samples from 5,000 to 14,000 generations. Standard assays and analysis of peak intensities using a Qiagen Pyromark 24 instrument were used to estimate the ratios of alternative alleles involving single-base substitutions in the cases of *rpsA* and *yghj* and a 16-bp deletion in the case of *gltB.* Again, control mixtures of REL607 and REL8953A cultures were used to verify the accuracy of inferred allele frequencies, including the correction for the relative DNA content in each strain as described above.

### Competitions to Test for Frequency-Dependent Interactions

The ancestral strain REL606 and the cells from population Ara-1 cannot grow on the sugar L-arabinose owing to a point mutation in the *araA* gene (Studier *et al.* 2009). However, one can obtain Ara^+^ mutants by inoculating large numbers of cells onto minimal arabinose (MA) plates; most of the resulting mutants are selectively neutral under the conditions of the LTEE, and the Ara phenotype serves as a useful marker in competition experiments used to measure relative fitness (Lenski *et al.* 1991). Thus, we isolated Ara^+^ mutants of certain evolved clones of interest by inoculating those clones onto MA plates. We then confirmed the neutrality of those mutations with competition assays involving a 1:1 volumetric ratio of the Ara^+^ mutants and their Ara-parents. These Ara^+^ mutants and their Ara-parents were then used to test for frequency-dependent interactions per the following procedure; this procedure is identical to that used to test for neutral Ara^+^ mutants, with the exception of using three volumetric ratios instead of one.

To begin a competition assay, two clones (one Ara^-^ and the other Ara^+^) were inoculated separately from freezer stocks into flasks containing LB medium and grown for 24 h at 37C. These cultures were then diluted 100-fold into saline solution, and 0.1 ml was transferred into 9.9 ml of DM25 (i.e., the same medium as used in the LTEE), where the cultures were again incubated for 24 h at 37C. This step ensured that the competitors were physiologically acclimated to the same conditions where they would compete.

The two clones were then mixed at three initial volumetric ratios (1:9, 1:1, and 9:1) and a combined 0.1 ml was added to 9.9 ml of DM25. An initial sample was taken immediately from this mixture and plated on TA agar (where Ara^-^ and Ara^+^ cells produce red and white colonies, respectively) to estimate the initial number of each type (*N*_i_). The mixture was then propagated for six days with daily 1:100 dilutions in DM25 medium. At the end of the experiment, a second sample was plated onto TA agar to estimate the final number of each type of cell (*N*_f_). The Malthusian parameter (*m*) was then calculated as the realized growth rate of a competitor over the competition, as follows: *m* = ln(100^*t*^ × *N*_f_/*N*-i)/*t*, where *t* is the number of days of dilution and re-growth, and where cell numbers reflect equivalent end-of-cycle values based on the dilutions used for plating. The fitness of one clone relative to the other was then calculated simply as the ratio of their Malthusian parameters.

For each pair of competitors, we performed 12 replicate assays at each initial ratio. The plate counts, fitness calculations, and R scripts used to analyze and plot the data are available at www.github.com/rohanmaddamsetti/Ara-l_Frequency_Dependence.

## RESULTS AND DISCUSSION

### Muller Plot of Allele Dynamics

Fig. 1a shows the dynamics across 20,000 generations of 42 spontaneous mutations that arose and reached detectable frequencies in the Ara-1 population of the LTEE. Each mutation is identified by the name of the gene in which it occurred (e.g., *topA*) or, for mutations in intergenic regions, by the adjacent genes (e.g., *yedW-yedX).* In a few cases, a numeral follows the name of a gene in order to distinguish mutations that affected the same gene (e.g., several *pykF* alleles). Twenty-five of these 42 mutations - all of them with labels preceded by dots (e.g., *topA* at the left and *rpsD* on the right) - eventually reached fixation (or nearly so) in the population. In other words, these mutations were on the line of descent leading to fully sequenced clones from generations 30,000 and 40,000. The other 17 mutations were transient (e.g., *fabR* on the left). That is, these alleles reached detectable frequencies - and in a few cases even achieved majority status (e.g., *acs-nrfA* between generations 8,500 and 12,500) - but they later went extinct. Trajectories for each mutation are also shown separately in Fig. S1.

### Not All Mutations Were Tracked

Fig. 1a contains a great deal of data, but when interpreting these dynamics it is important to realize that some information is missing because only a subset of all mutations were targeted in our genotyping assays. Many alleles were undoubtedly rare and transient; if an allele was not moderately abundant (at least several percent) in one of the generational samples that were deeply sequenced (Barrick and Lenski 2009), and if it was not on the line of descent leading to the clones sequenced in later generations (Barrick *et al.* 2009), then we could not design a targeted genotyping assay to detect it. Moreover, even some alleles that were known to fix in the population could not be accurately quantified using our genotyping method, (e.g., new transposon insertions and other structural mutations) and so information for those mutations is also missing from Fig. 1a. We assayed all 37 mutations in the 20,000-generation clone, REL8593A, that were on the line of descent leading to clones sequenced at 30,000 and 40,000 generations (Barrick *et al.* 2009). Twelve of these assays failed: six of the failures were structural mutations mediated by insertion-sequence elements, one was a single base pair insertion, and five were SNPs.

### Pure Drift Cannot Explain Evolutionary Rate

Random genetic drift alone cannot explain the large number of fixations observed or, for that matter, even one fixation. Each LTEE population began from a single haploid cell, and therefore any variant arose as a new mutation with an initial frequency of 1 */N,* where *N* is the population size. For a neutral mutation that eventually fixes in a population, the expected time to fix by random drift alone (i.e., without hitchhiking) is on the order of *N* generations (Kimura 1983). In the LTEE, *N* fluctuates between ~5 × 10^6^ and ~5 × 10^8^ as a consequence of the daily dilutions and regrowth (Lenski *et al.* 1991). These numbers are far too large to allow mutations to fix by pure drift in 20,000 generations, much less within the many fewer generations observed for the earliest fixations.

### Beneficial Drivers

We also know that mean fitness increased over the course of the LTEE. It increased in an almost step-like manner over the first few thousand generations (Lenski *et al.* 1991; Lenski and Travisano 1994), implying a series of fixations of beneficial driver mutations under positive selection, and it has continued to increase more gradually during the subsequent tens of thousands of generations (De Visser and Lenski 2002; Wiser *et al.* 2013). We also know or infer that particular mutations were beneficial because isogenic strains were constructed to measure the fitness effects of single evolved mutations, because similar mutations affecting the same gene reached high frequencies or fixed in many replicate LTEE populations, or both. These known or suspected beneficial drivers include 15 of the mutations in Fig. 1a including *topA* (Crozat *et al.* 2005, 2010; Woods *et al.* 2011), *fabR* (Woods *et al.* 2011; Deatherage *et al.* 2015), *spoT* (Cooper *et al.* 2003; Woods *et al.* 2011), three in *pykF* (Woods *et al.* 2006, 2011; Barrick *et al.* 2009), *mrdB* (Woods *et al.* 2006), *mrdA* (Woods *et al.* 2006), *malT* (Pelosi *et al.* 2006), *infB* (Barrick *et al.* 2009), *fis* (Crozat *et al.* 2005), *nadR* (Woods *et al.* 2006; Barrick *et al.* 2009), *pcnB* (Barrick *et al.* 2009), *iclR* (Barrick and Lenski 2009), and *rpsD* (Barrick *et al.* 2009). It is likely that additional mutations in Fig. 1a were also beneficial drivers, while some others may have been neutral or weakly deleterious hitchhikers that achieved transiently high frequency or fixation by virtue of beneficial mutations present in the same genomes.

### Clonal Interference

Clonal interference refers to the effect of competition between beneficial mutations that occur in different lineages in the same asexual population; its effects are clearly seen in Fig. 1a as wedges that open, expand, and then close. Owing to the absence of genetic exchange, two or more beneficial mutations that arise contemporaneously, but in different lineages, cannot be combined in the same genome. Instead, one will eventually prevail and the others must go extinct. The winning lineage might prevail because its initial driver mutation is more beneficial than the initial drivers in the contending lineages. However the winner might also depend on the effects of later beneficial mutations that arise in one or more of the contending lineages. Such an outcome is especially likely if the initial drivers in the different lineages have similar fitness effects, because that similarity means it would take longer for one lineage to exclude the other, thus providing more time for later beneficial mutations to occur and affect the outcome.

One interference wedge is seen near the very start as a mutation in *fabR* rises in frequency over the first 1000 generations, but that lineage declines precipitously during the next 500 generations before disappearing entirely (Fig. 1a). Five more wedges rise and fall between generations 1000 and 2500. Strikingly, three of them involve different mutations in one gene, *pykF.* Yet another mutation in *pykF* later went to fixation in this population (Schneider *et al.* 2000, Barrick *et al.* 2009), but it is not shown here because the type of mutation - an insertion of an IS *150* element - was not amenable to the method for detecting genetic polymorphisms used in this study. Moreover, different *pykF* mutations fixed in all 12 LTEE populations (Woods *et al.* 2006). Thus, there were numerous parallel increases of *pykF* mutations, both within and across populations, and this parallel evolution provides strong evidence that these were beneficial mutations.

Some additional interference wedges are also present later in the Muller plot including, most notably, an extremely large one that begins with a mutation in *rpsA* around generation 5000. This lineage became numerically dominant by generation 8000; it remained the majority for thousands of generations before a precipitous decline and sharp recovery between ~11,000 and 13,000 generations; and it finally petered out to extinction by generation 15,000. We will return to this episode in a later section on “Evidence for Frequency Dependence”.

### Rapid Increases in Mutation Frequencies

Except for the period from 5000 to 15,000 generations, most mutations that fixed in the Ara-1 population did so very quickly, usually within a few thousand generations. Although these dynamics are visually striking, they are not unexpectedly fast. Three issues come into play when we consider these dynamics. First, the alleles tracked in the Muller plot arose by mutation long before they are shown as being present. Only alleles with frequencies well above 1% could be reliably detected using our methods, but when any new mutation occurred its frequency was *1/N,* a mere 0.0000002% to 0.00002% depending on when the mutational event happened in the transfer cycle. In any case, each successful allele had to reach at least 0.00002% after surviving the first transfer event, which took place within the first day after it arose. Thus, mutations were hidden from view until they had increased in frequency by 50,000-fold or more.

Second, the fitness effects of beneficial mutations that evolved in the LTEE, and which have been measured by constructing and competing isogenic strains, are typically between ~1% and ~10% depending on the mutation (Barrick *et al.* 2009). Assuming a constant fitness difference, the logarithm of the ratio of the beneficial mutant to its progenitor should increase linearly (Dykhuizen and Hartl 1983). For a mutation that confers a 10% advantage, it takes only ~33 generations (doublings, as used in the LTEE) to rise from 1:100 to 1:10, another 33 generations to increase from 1:10 to 1:1 (i.e., 50%), another to go from 1:1 to 10:1, and one more to achieve near-fixation at 100:1. For this mutation, the entire ‘visible’ process would occur in ~ 130 generations; however, the time steps in Fig. 1a are 500 generations each, and so the fixation would likely be manifest in a single interval. For a mutation that gives a 1% benefit, each order-of-magnitude change in the ratio would take 10-fold longer, or ~330 generations. In that case, the visible fixation process would require ~1300 generations, which still falls within a mere three time steps in the Muller plot.

Third, in many of the fixation events, the initial rate of increase in a lineage was much steeper than the final rate. For example, the lineage with the *ybaL* allele went from undetectable in the 2000-generation sample to being the vast majority by generation 2500, but it did not fix until after generation 5000. Stated the other way, many lineages are unexpectedly persistent, such as the lineage that carried the *spoT* allele but lacked th *eyegl* and *ybaL* mutations, which hung on at low frequency from 2500 to 5000 generations. We can posit two explanations for this effect. First, as a consequence of clonal interference, a lineage that has a new beneficial mutation is initially competing primarily with its progenitor, but over time that lineage will increasingly compete against other lineages with different beneficial alleles, thereby slowing its ascent (Lang *et al.* 2011).

A second possibility is frequency-dependent selection, such that the unexpectedly persistent lineage has acquired some mutation that gives it a fitness advantage when it is rare; for example, it may more efficiently use a metabolic byproduct of the other lineage. This advantage when rare allows the lineage to survive longer than it otherwise could; in the absence of on-going evolution, its advantage when rare would allow it to persist forever. However, in this scenario the new majority lineage evolves, sooner or later, additional beneficial mutations that overcome the minority type’s advantage when rare, thus completing the extinction of the once-dominant lineage by the successor lineage. Incidentally, a somewhat different scenario occurred in another of the LTEE populations. In that population, two lineages stably coexisted by frequency-dependent selection. Both continued to fix beneficial mutations, and their relative abundance fluctuated dramatically over time. However, neither gained a sufficient advantage to drive the other extinct, and they have coexisted now for tens of thousands of generations (Rozen and Lenski 2000; Le Gac *et al.* 2012).

### Nested Fixations and Cohorts that Fix Together

In the Muller plot (Fig. 1a), we see many examples of nested fixations, in which one mutation begins its sweep to fixation in the background of a prior mutation that has not yet completed its own sweep to fixation. For example, before the *topA* allele has fixed, the *spoT* mutation has begun its rise; similarly, the *ybaL* mutation begins to spread before the *yegI* mutation has fixed. These nested fixations are not surprising; like clonal interference, they imply that mutations with fairly large beneficial effects are sufficiently common, given the population size, that more than one highly beneficial mutation is often present in contending subpopulations at the same time.

In other cases, however, we see what appear to be simultaneous fixations of two or more mutant alleles - what have been called ‘cohorts’ (Lang *et al.* 2013). One conspicuous example involved four mutations in the *mrdA, malT, nagC,* and *infB* genes that rose together starting at ~3,000 generations in the background carrying the *ybaL* mutant allele. Another case started around 12,500 generations with three mutations in the *pcnB, arcB,* and *ebgR* genes that spread and fixed together.

These quasi-simultaneous fixations may, at first glance, seem surprising. In fact, however, there are several plausible explanations for their occurrence, as explained below. An important consideration is that most of the time these mutant alleles spent rising in frequency occurred before they reached a frequency at which they could be detected; thus, the relevant dynamics for determining the order in which the mutations happened were hidden from our view. Also, the hypotheses below are not mutually exclusive; instead, two or more of the processes might be involved in any given cohort fixation.

*Simultaneous mutations.* The mutations could have occurred simultaneously; that is, the two or more changes in the DNA sequence might have taken place in the same replicating cell. However, given what we know about typical mutation rates in bacteria, in general, and the LTEE, in particular (Wielgoss *et al.* 2011, 2013), this explanation seems very unlikely. It is relevant to note that the Ara-1 population did not evolve a hypermutator phenotype during the 20,000 generations studied here, although it did so later (Barrick *et al.* 2009). It is also relevant that most mutations we were able to track using our methods were simple point mutations, rather than mutations involving mobile elements or other genetic mechanisms that may occur at much higher rates (Moxon *et al.* 1994; Cooper *et al.* 2001).

*Classic hitchhiking.* Neutral or even slightly deleterious mutations can fix in a population if they are physically linked to a beneficial mutation that goes to fixation. There is no horizontal gene exchange in the LTEE (Lenski *et al.* 1991; Lenski 2004), and so the LTEE populations have perfect linkage. If the beneficial mutation that is eventually fixed occurs in a background that already has the neutral or deleterious mutation, then the hitchhiker and beneficial driver will fix at the same time. If the order is reversed, such that the neutral or slightly deleterious mutation occurs in the background of the beneficial mutation, but one or more generations later, then the situation becomes more complex. If the hitchhiker is deleterious, then it may rise in frequency for a while but will not fix, because the lineage that carries the beneficial mutation alone will out-compete the lineage with both mutations. If the hitchhiker is neutral, then the double mutant will rise in parallel with the lineage that carries only the beneficial driver, but the hitchhiker should not fix. For example, if a neutral hitchhiker occurs when the lineage with the beneficial mutation has expanded to four cells, then we would expect the double mutant to reach a frequency of ~25% when the beneficial mutation fixes in the population. In both cases, these scenarios are simplifications because they ignore the effects of clonal interference and random drift. The latter is important especially while the beneficial allele is still very rare. In particular, although a hitchhiker might occur after the beneficial driver, if the genotype that contains only the beneficial mutation was lost by drift (e.g., during a transfer soon after it appeared), then the hitchhiker and driver mutations would fix simultaneously.

*Beneficial co-drivers.* The term hitchhiker is usually applied only to neutral or deleterious mutations, as discussed in the preceding paragraph. However, two or more beneficial mutations that occur in the same lineage can help drive one another to fixation, including cases where one or both mutations might not be able to fix by themselves owing to clonal interference from single beneficial mutations of larger effect (Cooper *et al.* 2001; Schiffels *et al.* 2011). In essence, such mutations are co-drivers that receive a boost from their partner, similar to the boost that a hitchhiker would get. Although the beneficial mutations may occur sequentially and, in fact, many generations apart, they may nonetheless appear to fix simultaneously. The reason for the apparent simultaneity of fixation hinges on the dynamics of selection coupled with the limits of detection of rare genotypes.

Consider a lineage bearing a mutation that confers a 5% fitness advantage, and which survived extinction by random drift. Using the same framework as in the section on “Rapid Increases in Mutation Frequencies,” the ratio of that lineage to its progenitor should increase by an order of magnitude in ~66 generations. After 132 generations, ~100 cells will have the beneficial mutation. Now imagine that, at this point, a second beneficial mutation occurs in the background with the first mutation, and that the double mutant has a 5% advantage relative to the single mutant and a 10% advantage over the numerically dominant type without either mutation. Again, we will assume that the double mutant escapes extinction by drift, because we are only interested in those cases that leave a record of fixations. The double mutant should increase by an order of magnitude relative to the single mutant in ~66 generations, and relative to the non-mutant progenitor in 33 generations. Let us now assume that the mutant alleles can be detected only when they become 1% of the total population. For purposes of these calculations, we will use 107 as the population size; this value is conservative because the effective population size, taking into account the daily transfer cycle in the LTEE, is somewhat larger (Lenski *et al.* 1991). Absent the second beneficial mutation, the single-mutant lineage would require ~330 generations to increase five orders of magnitude and reach the 1% frequency where it could be detected. Although the double mutant appeared 132 generations later, it requires only ~165 generations to reach the 1% threshold in the population, which remains numerically dominated by the genotype with neither beneficial mutation. Then, after 99 generations more—or 396 generations since the first mutation and 264 since the second— the double mutant would be ~100-fold more common than the single mutant. With the 1% limit of resolution, and with samples taken at 500-generation intervals, it would thus appear as though the two mutations had simultaneously fixed, even though they arose many generations apart and were, strictly speaking, nested.

*Positive epistasis.* This explanation is, in essence, an extension of the previous one. In the LTEE, there is an overall tendency toward diminishing-returns (i.e., negative) epistasis between beneficial mutations with respect to their fitness effects (Khan *et al.* 2013; Wiser *et al.* 2013), although there are also exceptions where mutations exhibit positive epistasis (Blount *et al.* 2012). Even if negative epistasis is more common, cases of positive epistasis may play a disproportionate role in cohort fixations. This effect can be easily understood by imagining a scenario like the one discussed in the previous paragraph, except now imagine that the two mutations together give a 15% benefit, whereas either alone would provide only a 5% benefit. In this case, the double mutant would overtake the single mutant that much sooner, obscuring the sequential order faster and more completely; it would do so even if the two mutations were more separated in time, if their individual beneficial effects were smaller, or both.

*Frequency-dependent selection.* Two lineages can stably coexist if, as a result of negative frequency dependent interactions, each type has a fitness advantage when it is rare. As long as the stable equilibrium persists, each lineage can accumulate an arbitrarily large number of distinguishing mutations by natural selection and hitchhiking (Rozen and Lenski 2000; Herron and Doebeli 2013). Now imagine that, at some later time, a beneficial mutation or combination of mutations arises in one lineage, the fitness effect of which is large enough to disrupt the stable equilibrium and drive the other lineage extinct. At that time, this hyper-beneficial mutation will not only fix but also drive to fixation in the total population all the other alleles that previously had fixed only in its lineage. In the next section, we will present evidence that such a scenario played out in population Ara-1.

### Evidence for Frequency Dependence

We now turn our attention to the unusual dynamics that occurred in this population between 5000 and 15,000 generations (Fig. 1a). To better resolve the dynamics, including especially the linkage relationships among various alleles, we genotyped 90 clonal isolates sampled from generations 7500 (Fig. 1b) and 10,000 (Fig. 1c) for the presence or absence of the various mutations. The earliest discernible events involved the simultaneous rise of two distinct clades from within the blue-colored background that bears a cohort of four mutations (including *mrdA*) that eventually fixed (Fig. 1a). One of the two clades began with a mutation in the *rpsA* gene; it is aqua-colored at first, then various shades of green as more mutations accumulated with time (Fig. 1a). The nested pattern of the first two mutations in *rpsA* and *acs-nrfA* was evident in the clones from generation 7500 (Fig. 1b), but by generation 10,000 these two mutations were fully concordant (Fig. 1c). Three more mutations - in *atoS, nuoG,* and *nuoM -* emerged as distinct sub-clades in the later sample, but they did not persist (Fig. 1c). The other main clade appeared first as a cohort of four mutations (including *gltB*); it is shown in chartreuse initially, with shades of yellow, red, pink, and purple used as more mutations arose (Fig. 1a). In the clones sampled at 7500 generations, it was impossible to discern the order of those first four mutations, although the next three mutations that arose in this lineage were already present as nested subsets at that point. By generation 10,000, six mutations in this clade had become fully concordant; two mutations that would eventually fix (*pflC* and *nadR-*2) were nested, as were four others that did not persist. Following their appearances after 5000 generations, both major clades increased in number, with the former one (harboring the *rpsA* mutation) becoming the clear majority by 8000 generations and remaining so for most of the next few thousand generations. It then experienced a precipitous decline, a sharp recovery, and another precipitous decline, followed by its extinction. We confirmed that these sharp fluctuations were not artifacts of our genotyping method by also using pyrosequencing to track three diagnostic alleles that distinguished the two clades (Fig. S2). By the end of this episode, at least 14 mutations (from *gltB* to *iclR,* and probably others not detected by our methods) had piled up on the line of descent before the lineage that ultimately prevailed finally drove the other clade extinct by 15,000 generations.

*Clonal interference alone is insufficient.* One might be tempted, at first glance, to suggest that this episode was simply a protracted case of clonal interference. We know that the rate of fitness increase decelerated over time (Wiser *et al.* 2013) and that the beneficial fitness effects of mutations that fixed later tend to be smaller than those that fixed earlier owing to diminishing-returns epistasis (Khan *et al.* 2011). Together, these facts imply that later fixations usually took longer than earlier ones, thereby providing the opportunity for more drawn-out bouts of clonal interference. However, this explanation does not account for two important features of the data: (i) there were rapid fixations after, as well as before, this episode; and (ii) sharp reversals in the relative abundance of the two lineages occurred within this episode. Moreover, as explained earlier, these dynamics are entirely consistent with the smaller selection coefficients that prevailed in the later generations. Therefore, we think that some other process than simple clonal interference must have contributed to this extremely drawn-out set of fixations.

*Hypothetical scenario with frequency-dependent fitness.* We hypothesized that a negative frequency-dependent interaction stabilized the relative abundance of the two clades, while beneficial mutations within each clade buffeted their abundances until, eventually, one drove the other extinct (Fig. 2). Under the null hypothesis, the relative fitness of members of the two clades, taken at the same point in time, does not depend on their relative abundance in the population (Fig. 2a). The alternative is that each type has an advantage when rare, such that there is a stable equilibrium (i.e., a relative fitness equal to unity) at some intermediate frequency (Fig. 2b). Now imagine that beneficial mutations occur in both lineages that alter the shape of the fitness function (Fig. 2c). We show these changes as affecting the intercept, but not the slope, of the fitness function, as though the mutations provide a general benefit that does not affect the frequency-dependent interaction; however, one can imagine similar scenarios where the slope changes. In the scenario illustrated, the initial equilibrium frequency of clade C2 is ~0.2. A beneficial mutation in clade C1 drives the equilibrium frequency of C2 down to ~0.05 (green line), and then a later beneficial mutation in C2 drives its equilibrium frequency up to ~0.5 (blue line); finally, though, another mutation in C1 pushes the equilibrium frequency of C2 into negative territory (red line) and C2 then goes extinct, assuming it gets no further beneficial mutations, because it is less fit than C1 at all frequencies (Fig. 2c).

**Figure 2.**
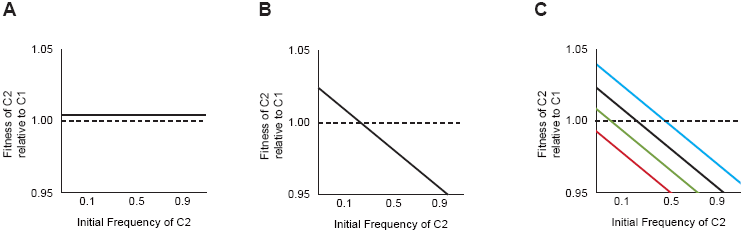
Hypothetical scenario showing the interplay between negative frequency dependence and ongoing beneficial mutations. (A) Null case, in which the fitness of clade C2 relative to clade C1 is independent of their relative frequencies. (B) Static negative frequency dependence, where the fitness of C2 relative to C1 declines as the frequency of C2 in the population increases. (C) Dynamic frequency dependence, where ongoing mutations in each clade provide generic fitness benefits. Although the mutations do not affect the frequency-dependent interaction per se, they affect the location and existence of the stable equilibrium. In the example shown here, the first beneficial mutation (green line) occurs in Cl and lowers the equilibrium frequency of C2; the second beneficial mutation (blue line) occurs in C2 and raises its equilibrium frequency; and a third beneficial mutation (red line) occurs in C1 that eliminates the equilibrium and drives C2 extinct.

*Experimental tests of frequency dependence.* We ran competition experiments to test whether representatives of the two lineages exhibited frequency-dependent interactions at two time points. We chose clones from the 7500-generation sample that represented the most derived genotypes within clades C1 and C2; they differed by at least nine mutations (Fig. 1b). They competed head-to-head starting at three different ratios, with 12-fold replication for each ratio. We calculated their relative fitness as the ratio of their realized growth rates during the competition assay. We saw no evidence of frequency dependence (Fig. 3a); of course, we cannot exclude the possibility of a very small effect, on the order of 1% or less. It is also noteworthy that there was no detectable difference in fitness between the two competitors, even though they differed by so many mutations and both were increasing in abundance relative to the overall population in which they arose (Fig. 1).

**Figure 3.**
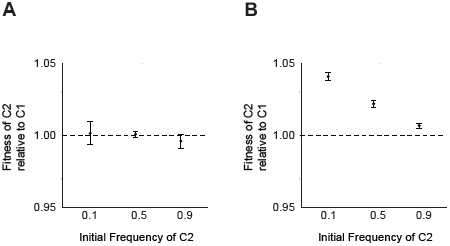
Changing nature of interaction between clones from two clades over time. (A) Clonal isolates from clades C1 and C2 sampled from population Ara-1 at 7500 generations show no frequency-dependent interaction. Although they differ by at least 9 mutations, there is no significant difference in their fitness at any of the three initial ratios tested. (B) Clonal isolates from the same clades at 10,000 generations show a strong negative frequency-dependent interaction, with the fitness of the C2 clone declining as its frequency increases. The C2 clone has a significant advantage at all of the initial ratios tested, although there may be a stable equilibrium with C2 at a frequency above those tested. Although C2 eventually drove C1 extinct, ongoing beneficial mutations evidently affected the dynamics (Fig. 1A). Error bars show 95% confidence intervals.

We repeated this experiment except using the most derived genotypes from clades C1 and C2 at 10,000 generations. In particular, the C1 clone had a mutation in *nuoM,* and the C2 clone had a mutation in *nadR* that was the latest mutation that subsequently fixed in the Ara-1 population. In this case, we observed very strong frequency dependence; the C2 clone had a fitness advantage of 5.4% when its starting frequency was 10%, an advantage of 2.9% when its initial frequency was 50%, and an advantage of 0.9% when it started at 90% (Fig. 3b). Although the C2 clone was fitter than the C1 clone at all three initial ratios tested, the trend suggests the possibility of stable coexistence with C1 at an equilibrium frequency of <5%. Note, however, that this hypothetical equilibrium frequency does not match the frequency of clade C1 observed in the 10,000-generation sample, when it was roughly half the total population (Fig. 1). This discrepancy is not surprising, though, because the competitions involved single representatives from two clades that were continuing to evolve and adapt over time. The important point to take away from the frequency-dependent interaction is this: as clade C2 spread, its advantage over C1 diminished. The frequency dependence thus helps to explain, at least in part, why this selective sweep took so much longer, and involved many more mutations, than other sweeps that both preceded and followed it.

*Failed speciation.* Defining speciation is often problematic, and especially so for asexual organisms like those in the LTEE. Nonetheless, one interesting way to think about speciation in bacteria posits that it occurs when lineages have diverged to occupy ecological niches that are sufficiently different that selective sweeps can happen in one lineage without driving the other extinct (Cohan 2006; Cohan and Perry 2007). This model fits with the general idea that species are cohesive groups that have diverged irreversibly (Templeton 1989). The importance of reproductive isolation between sexual species is that it locks in the requisite divergence, but asexual organisms can also diverge irreversibly by adapting to different ecological opportunities (Cohan 2002). Of course, any definition that depends on a multistep temporal process is likely to involve intermediate states - grey areas - of ambiguous status. The divergence and eventual fates of the clades C1 and C2 illustrate that ambiguity. That is, these two lineages diverged ecologically and accumulated different beneficial mutations over several thousand generations, and one can imagine that they might have coexisted indefinitely. However, despite the ecological divergence that led to their negative frequency-dependent interaction, the two clades continued to compete and their fates remained entangled. In time, the further adaptation of one clade caused the community of two nascent species to collapse to a mono-specific population. Without the “fossil record” of frozen samples, studies of later generations would reveal nothing of this incipient, but ultimately failed, process of speciation.

### Conclusions

We tracked the dynamics of several dozen mutations in an *E. coli* population as it evolved in and adapted to a simple laboratory environment for 20,000 generations. The population started from a single haploid cell, and the bacteria in the experiment lack any mechanism for horizontal gene transfer. Glucose was the limiting resource, and its concentration in the culture medium was set low to reduce the cell density and thus, it was thought, simplify the evolutionary dynamics in two respects (Lenski 2004). First, a low population density should reduce the concentration of metabolic byproducts and thereby limit the opportunity for frequency-dependent interactions. Second, a low population size should reduce the rate at which beneficial mutations arise and thereby reduce the impact of clonal interference.

Despite the simplicity of these conditions, the dynamics are rich and complex. As expected, we observed many selective sweeps in which derived alleles replace their ancestral counterparts. The speed with which the sweeps occurred is inconsistent with pure genetic drift, but consistent with what is known about the fitness effects of individual mutations in this experiment, given also the limits of detection for the mutations. We also observed, as expected, many cases of clonal interference. We undoubtedly documented far fewer instances of interference than actually occurred, because we tracked only mutations that were known to have either fixed in the population or been present at substantial frequency in a few generational samples that were previously screened for polymorphisms (Barrick and Lenski 2009).

Two other features of the genome dynamics were equally conspicuous but more surprising, especially given the simplicity of the experimental conditions. One such feature was the drawn out and temporally complex interaction of two clades, which required several thousand generations before one clade eventually excluded the other. This episode reflected a combination of frequency-dependent selection and the rise of new beneficial mutations in both clades that buffeted their relative numbers before one clade finally gained an insurmountable advantage. Previous work has revealed an even longer-lasting coexistence of two clades in another of the LTEE populations (Rozen and Lenski 2000; Le Gac *et al.* 2013). In that population, beneficial mutations also buffeted the abundances of the clades, but the frequency dependence was sufficiently strong that they have continued to coexist for several tens of thousands of generations (Le Gac *et al.* 2013). Our study, along with the work on that other population, shows that frequency-dependent interactions emerged in the LTEE, despite the low resource concentration intended to limit their importance (Elena and Lenski 1997). Our study also shows that such interactions can be transient and might appear, at least superficially, to be cases of very drawn out clonal interference.

The second unexpected feature of the genome dynamics that we observed is the prevalence of cases in which two or more mutations appear to have fixed more or less simultaneously in the population. These cohorts seem to be at odds with the expectation that fixations should be sequential or nested (i.e., one mutation arising in the background that contains the other before it reaches fixation). In a recent study of yeast populations, Lang *et al.* (2013) found similar cohort fixations by deeply sequencing genomes at many time points. However, the explanation for these mutational cohorts remains unclear. We proposed several possibilities, ranging from the conceptually simple (but unlikely) idea that the mutations occurred simultaneously to more complicated scenarios that invoke epistasis and frequency dependence. We think the most parsimonious explanation - the null hypothesis from a population-genetic perspective - is that these cohorts are an illusion caused by the limited resolution of the actual genome dynamics. That is, mutations are only detected in these experiments after they have become fairly common - at least one percent - and by that time they have already increased in frequency by many orders of magnitude. As a consequence, given two beneficial mutations that occurred sequentially - but in sufficiently close temporal proximity - the double mutant may reach a detectable frequency before the first mutation alone or even when the first mutation alone never reaches a detectable frequency (especially given the 500-generation interval between successive samples in our data, and the 80-generation sampling interval in Lang *et al*.).

It is not obvious to us whether the different scenarios, including the null hypothesis, can be distinguished empirically given the limited resolution of population-genomic methods, the temporal gaps between population samples, and the very small fitness effects (including frequency-dependent and epistatic interactions) that are likely to be relevant to any particular case. Instead, we suggest that numerical simulations could provide an important next-step toward resolving the causes of the cohorts. By performing simulations, one can explore what rates of mutations and distributions of fitness effects would give rise to the appearance of cohorts for various allele detection limits and sampling schemes. The inferred rates and distributions could then be compared to corresponding rates and distributions estimated in other ways, such as from population mean-fitness trajectories (Wiser *et al.* 2013), to determine if they are compatible. Whatever the findings from these theoretical studies might be, the results of our study and the rapidly growing body of data from the LTEE and other evolution experiments show the exciting challenges that remain for describing and understanding the dynamics of genomic and phenotypic evolution in microbial populations, even under seemingly simple conditions.

## Acknowledgments

We thank Zachary Blount and Noah Ribeck for discussions; Steven Valenziano for help with producing figures; Jeff Landgraf and Cecil Harkey for assistance with the genotyping and pyrosequencing assays; Jeff Morris, Magdalena Felczak, and Louis King for protocols and help with the flow cytometry; and Neerja Hajela for assistance in the laboratory. This work was supported by the National Science Foundation (DEB-1019989 to R.E.L.), the National Institutes of Health (R00-GM087550 to J.E.B.), the NSF BEACON Center for the Study of Evolution in Action (DBI-0939454), and a National Defense Science and Engineering Graduate Fellowship to R.M.

